# Scale-free dynamics in animal groups and brain networks

**DOI:** 10.1101/2020.12.02.409029

**Authors:** Tiago L. Ribeiro, Dante R. Chialvo, Dietmar Plenz

## Abstract

Collective phenomena fascinate by the emergence of order in systems composed of a myriad of small entities. They are ubiquitous in nature and can be found over a vast range of scales in physical and biological systems. Their key feature is the seemingly effortless emergence of adaptive collective behavior that cannot be trivially explained by the properties of the system’s individual components. This perspective focuses on recent insights into the similarities of correlations for two apparently disparate phenomena: flocking in animal groups and neuronal ensemble activity in the brain. We first will summarize findings on the spontaneous organization in bird flocks and macro-scale human brain activity utilizing correlation functions and insights from critical dynamics. We then will discuss recent experimental findings that apply these approaches to the collective response of neurons to visual and motor processing, i.e. to local perturbations of neuronal networks at the meso- and microscale. We show how scale-free correlation functions capture the collective organization of neuronal avalanches in evoked neuronal populations in nonhuman primates and between neurons during visual processing in rodents. These experimental findings suggest that the coherent collective neural activity observed at scales much larger than the length of the direct neuronal interactions is demonstrative of a phase transition. We discuss the experimental support for either discontinuous or continuous phase transitions. We conclude that at or near a phase-transition neuronal information can propagate in the brain with the same efficiency as proposed to occur in the collective adaptive response observed in some animal groups.

## Introduction

The collective movement of animal groups has been the subject of great interest for many decades, with the early work focusing on model simulations (Aoki, 1982;Reynolds, 1987). It is now well accepted that collective properties in animal groups are closely related to the general study of collective phenomena in physics, which initially was focused on phase transitions in equilibrium systems composed of many, locally interacting particles (Stanley, 1971;Ma, 1976;1985), but eventually was expanded to include far-from-equilibrium systems (Meakin, 1987 ;Kertesz and Wolf, 1989;Martys et al., 1991). Many biological systems were found to fit into this latter category specifically when considering systems of self-driven particles to model movements of ants (Millonas, 1992;Rauch et al., 1995), fish schools (Huth and Wissel, 1992) and bird flocks resulting in the seminal model by Vicsek and colleagues for flocking in biological systems based on local interactions impacted by noise (Vicsek et al., 1995). Since then, variations of the Vicsek model (Gregoire and Chate, 2004;Chate et al., 2008) as well as other models that utilize attraction and distance rules (Couzin et al., 2002;Romanczuk et al., 2009) have been combined with experimental observations to capture population dynamics of many species such as locust swarms (Huepe et al., 2011), ants (Gelblum et al., 2016), fish schools (Tunstrøm et al., 2013), migrating white storks (Nagy et al., 2018) and cycling pelotons (Belden et al., 2019) with a major goal to understand the emergence of collective behavior from the mechanistic interactions between individuals (for a review, see e.g. (Wang and Lu, 2019)).

These observations support the idea that biological systems seem to be naturally poised near a phase transition (Bak, 1996), where they might benefit from order yet maintain adaptability to changing environmental conditions, an idea that is increasingly gaining attraction including the brain (Chialvo, 2010;Mora and Bialek, 2011;Plenz, 2012;Hesse and Gross, 2014;Plenz and Niebur, 2014). The initial theoretical debate has been enriched recently by an ever-improving ability to simultaneously track many biological elements (neurons, birds, midgets, etc.) over time, such that now the ideas are being challenged and contrasted by the experimental findings in the usual manner of statistical mechanics.

In this note, we focus on the behavior of the system correlation properties, the central tenet of statistical mechanics. For the sake of discussion, our starting point will be the work by Cavagna and colleagues in 2010, who demonstrated that starlings in a flock exhibit spatial correlations much longer than the length of direct interactions between neighboring birds (Cavagna et al., 2010;Cavagna et al., 2018). Specifically, they showed that the correlation length, i.e., the distance at which correlations drop below zero, grows monotonically with flock size (Fig. 1A) and is, therefore, scale-free. The absence of any characteristic scale in the correlations is known to be a hallmark of critical systems (Wilson, 1979). For the human brain, early evidence of scale-free correlation functions was found for ongoing neuronal activity assessed indirectly using the blood oxygen level dependent signal (BOLD) (Expert et al., 2011) followed by the demonstration of correlation length to grow with the size of the observed brain region (Fig. 1B; (Fraiman and Chialvo, 2012)), exactly as was described for starling flocks. These remarkable population-spanning correlations were replicated for a network model of the brain with experimentally based interareal connectivity when the network dynamics was tuned to criticality (Haimovici et al., 2013). Since then, they have also been observed for bacterial colonies (Chen et al., 2012), insect swarms (Attanasi et al., 2014b) and globular proteins (Tang et al., 2017;Tang et al., 2020). Here, we explore specifically the analogy in scale-free correlations between animal groups and brain dynamics at the scale of local population activity during motor outputs in nonhuman primates and down to the cellular scale of single neuron interactions during sensory processing in mice. We will demonstrate that this analogy goes beyond phenomenology and shares the same formal scaling relations which suggest common underlying principles.

**Figure 1:**
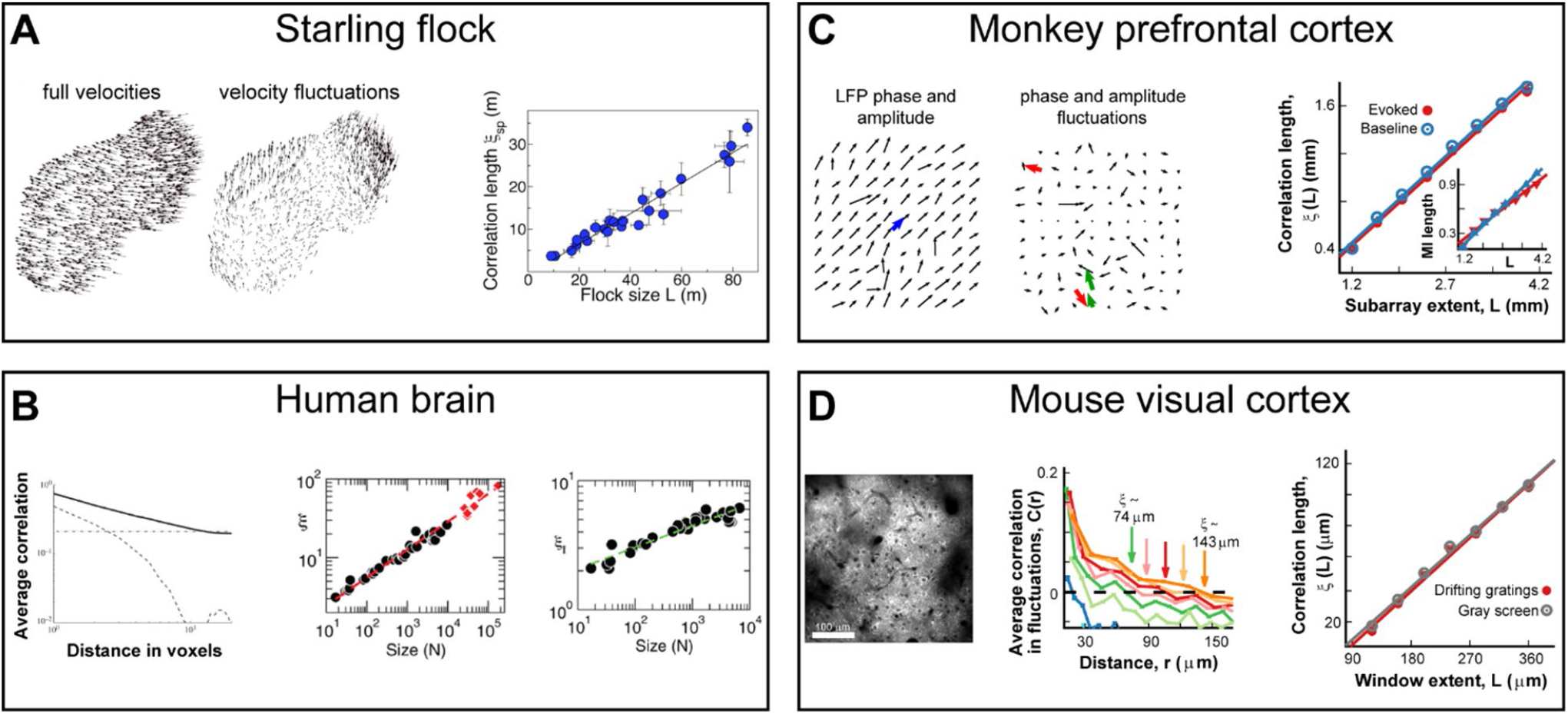
Scale-free growths in correlations length is observed in bird flocks and the mammalian brain at different scales and using different recording techniques. **(A)** Correlations in the velocity fluctuations of pairs of starlings in flocks of different sizes. Fluctuations are obtained by subtracting from each bird’s velocity vector (*left*) the center-of-mass velocity of the flock (*middle*). Correlation length, defined as the distance at which correlations of the fluctuations reaches zero, scales linearly with flock size (*right*) in line with expectations from critical dynamics. Adapted from (Cavagna et al., 2010). **(B)** Correlations obtained from blood oxygenated level dependent (BOLD) signals using fMRI to measure ongoing neuronal activity of the human brain. *Left*: Average correlation between voxel pairs drops with distance between voxels as a power law (*solid line*), while phantom data drops exponentially (*dashed line*) and spatially shuffled data is constant (*dotted line*). Adapted from (Expert et al., 2011) *Middle*: Correlation length, ξ, from fluctuations in BOLD data scales linearly with the size of the brain area observed (*black circles*) or when pooling areas together (*red diamonds*). Adapted from (Fraiman and Chialvo, 2012). *Right*: Mutual information between voxel pairs decays with pair distance, allowing for the definition of “mutual information length”, ξ_I_, in analogy to correlation length. ξ_I_ scales linearly with the size of the brain area observed (*black circles*). Adapted from (Fraiman and Chialvo, 2012). **(C)** Correlations in the fluctuations of LFP amplitudes from prefrontal cortex in nonhuman primates during a working-memory task using high-density microelectrode arrays. *Left/middle*: LFP vectors depicting phase and amplitude on the array without/with subtraction of the population average (*blue arrow, left*) in analogy to velocity distributions in flock data. *Right*: Correlation length scales linearly with (sub)array size for both ongoing (*blue*) and evoked (*red*) data. Adapted from (Ribeiro et al., 2020). *Inset*: Mutual information length scales linearly with (sub)array size for both ongoing (*blue*) and evoked (*red*) data. **(D)** Correlations in the fluctuations of neuronal activity from primary visual cortex in mice during visual stimulation using 2-photon imaging. *Left*: Example field-of-view showing cells used for the analysis. *Middle*: Average correlation of activity fluctuations between pairs of neurons decays with distance as well as with the size of the observed window (*colors*). *Right*: Correlation length scales linearly with observed window size for both gray screen (*gray*) or drifting gradings (*red*). Adapted from (Ribeiro et al., 2020).

### Scale-free correlations in response to external perturbations

The absence of a central control for the emergence of order lies at the heart of collective phenomena. With respect to animal groups this remarkable feature is also known as “coordination” and allows animals to stay together for protection in the face of predators (Powell, 1985;Terborgh, 1990;Krause and Ruxton, 2002) or to enhance foraging (Krebs, 1973;Munn and Terborgh, 1979;Greenberg, 2000). This collective response thus requires information about a local predator or local food source to be translated into a coordinated flock response for escape behavior or foraging to be successful. Several studies have now demonstrated how swarms can achieve such de-centralized coordination using local interactions between neighbors (Gregoire et al., 2003;Sumpter, 2006;Strombom, 2011;Bialek et al., 2012;Vicsek and Zafeiris, 2012;Ling et al., 2019b).

Predominantly local interactions are also characteristic for many brain networks, specifically as found for the cortex in mammals (Markram et al., 2015). Like a bird in a flock, the ‘action’ or output of a cortical neuron depends largely on the activity of its intracortical neighbors (Boucsein et al., 2011). The response to external perturbations of a flock, e.g., by the local intrusion of a predator, also invite interpretations similar to the response of a cortical network to external inputs. Those inputs directly affect only a small proportion of all neurons, e.g. through input from the thalamus (Bruno and Sakmann, 2006) or from other cortical regions, and thus are analogous to local perturbations of ongoing network dynamics (Arieli et al., 1996). And although neurons in a network do not change physical positions in relation to one another like birds, they may change their interaction neighborhood over time by strengthening or weakening their direct connections through synaptic plasticity. The mechanisms by which neuronal networks can propagate information quickly and flexibly to very distant, but not directly interacting, neurons are less clear though. Thus, inspired by the flock results we searched for evidence of scale-invariant correlations in brain activity in response to sensory input.

We recently explored the behavior of neuronal correlation functions at scales closer to direct neuronal interactions (Ribeiro et al., 2020). At the scale of a cortical area (i.e., the mesoscale of millimeters), we measured the distribution of the so-called local field potential (LFP) with high-density microelectrode arrays implanted in the premotor and prefrontal cortices of nonhuman primates performing a self-initiated motor task and a working memory task, respectively. The LFP extracts the local synchronization of neuronal groups and its emergence and propagation thus tracks the spatiotemporal evolution of population activity at a spatial resolution of several hundred micrometers with millisecond precision. At the scale of the cortical microcircuit (i.e., the microscale of microns), we measured the intracellular calcium dynamics in pyramidal cells expressing the genetically encoded calcium indicator YC2.6 in superficial layers of the primary visual cortex in awake mice passively viewing drifting gratings. The fluorescent indicator closely tracks the action potential firing in individual pyramidal neurons, which allows for a cellular reconstruction of spatiotemporal population activity with micrometer spatial resolution and sub-second temporal precision. At both scales, we observed the linear growth of the correlation length as a function of the linear size of the sampled area during sensory processing and motor output (Figs. 1C, D). Remarkably, these scale-free correlations were similarly present during rest and evoked responses from the sensory/motor stimulation (Figs. 1C, D; see also (Ribeiro et al., 2020)). In line with previous results for the whole brain (Fraiman and Chialvo, 2012), the mutual information found in neuronal activity also behaved in a scale-free manner. By measuring how the mutual information between pairs of electrodes decays with distance, we showed that the “mutual information length” grew linearly with system size, just like the correlation length, for ongoing and evoked neuronal at the mesoscale (Fig. 1C, inset).

Animal groups exhibit collective behavior in space during motion, in contrast to the brain, where activity propagates in high-dimensional networks and neurons themselves are stationary. These differences come into focus when considering scaling of correlation length by the spontaneous breaking of continuous rotational symmetry as is the case for orientation in space. In this case, global ordering can emerge in the absence of criticality at lower temperatures including the presence of powerlaw decay in space (Goldstone’s theorem; (Goldstone, 1961)). For this reason, Cavagna and colleagues (Cavagna et al., 2010) also investigated correlations in the speed of birds, for which that argument does not apply: whereas orientation could be seen as a soft mode (being bound, they have a “soft” degree of freedom), speed in principle is unbounded and thus is considered a so-called “stiff” mode. In the case of brain activity, the Goldstone’s theorem does not apply, at least for the data presented here, since there is no continuous symmetry that can be broken or soft modes. It needs to be noted that although a “pseudo” phase can be extracted from the LFP using a Hilbert transform of the original time series (Yu et al., 2017) the work of Ribeiro et al. (2020) used only the change in LFP amplitude (which is unbounded) to compute the correlation length. Furthermore, similar results were obtained when using binarized negative excursions of the LFP below a certain threshold (so-called nLFPs, which represent the local, synchronous firing of neurons around the electrode; see (Yu et al., 2017)), calcium traces or deconvolved spikes (Ribeiro et al., 2020), all of which are analogous to speed in animal movement.

### Interaction length versus correlation length

As commented, many animals living in groups synchronize their behavior to that of their neighbors. In that manner, they can spend less time on the lookout for predators and more time feeding or resting (Bednekoff and Lima, 1998). If animals were required to be on alert to the behavior of distant group members, more resources would need to be allocated to group observation. Obviously, this requirement might not even be possible, e.g., for herds that are confined to a plane where observation of distant members is obscured or for very large animal groups in general. The attention towards neighbors is accounted for by most models of collective behavior in animal groups, which, considering local interactions (Vicsek et al., 1995;Cucker and Smale, 2007;Wang and Lu, 2019), are able to capture the synchronization of animals to their neighbors as found for red deer (Rands et al., 2014) and recently for black-headed gulls (Evans et al., 2018). In a more extreme example, mosquitofish were shown to only respond to their single nearest neighbor (Herbert-Read et al., 2011). Thus, regardless of whether interactions between animals depend on metric or topological distance (Ballerini et al., 2008;Ginelli and Chate, 2010;Strandburg-Peshkin et al., 2013), it is probably safe to say that keeping track of nearby neighbors is a preferred behavioral strategy in groups. On the other hand, this strategy requires that information pertinent for the individual survival must travel efficiently throughout the entire group, independently of the group size. In physics, this feature of transforming local (short-range) interactions into global (long-range) correlations, is known to be present in systems (almost exclusively) at criticality (Wilson, 1979). Support for this concept comes from the work of Cavagna and colleagues who employed a maximum entropy approach to infer the effective interactions from individuals in a natural flock and showed that the interaction range decays exponentially over the range of just a few individuals (Cavagna et al., 2015). Additionally, Calvão & Brigatti’s model (Calvao and Brigatti, 2019), which is an implementation of the classical “selfish herd hypothesis” (Hamilton, 1971), is composed of local-interacting agents which collectively undergo a discontinuous phase transition. Their model successfully reproduces the behavior observed in nature for midge swarms including long-range correlations (Attanasi et al., 2014a;b).

For the brain, direct interactions between neurons exhibit a far more complex and selective organization than nearest neighbor relations. Neuronal interaction in the cortex includes a dominant number of direct short-range connections onto which long-range connections are superimposed that link distant cortical regions within and between hemispheres. Accordingly, the observation of long-range correlations might arise from short-range interactions at critical dynamics, from long-range connections independent of dynamical regimes, or both. To disambiguate this, we have simulated critical dynamics in a neuronal network with a precisely defined characteristic size for its connections and evaluated how the correlation function changes for distances beyond the short interaction range. We found that there is a clear change in the behavior of the correlation function at the interaction range, with correlations growing much faster for distances up to this point, confirming our experimental findings in primary visual cortex. The obtained interaction distance was similar to the characteristic distance at which two pyramidal cells in layers II/III are connected anatomically (Levy and Reyes, 2012;Seeman et al., 2018). These results suggest critical dynamics in combination with short interactions to be a major factor behind the observed correlation length scaling at the microscale and indirectly suggest that as in the case of animal flocks, the information about a local input or perturbation can rapidly propagate through the entire system.

### Effects of the heterogeneity of the elements on the correlation structure

Although some early works have taken heterogeneity and self-sorting into account (Couzin et al., 2002), animal group behavior has been mostly studied assuming homogeneous behavior of the individual (Ero et al., 2018;Gouwens et al., 2019). More recently, the effects of heterogeneity within groups has gained increased attention (for a review, see e.g. (King et al., 2018)). For instance, it has been shown that body size affects the strength of social interactions and the spatial organization of fish schools (Romenskyy et al., 2017). For jackdaws, a bird species that form lifelong pair-bonds, social relationships between different birds lead to the appearance of sub-structures within a flock. Pair-bonded jackdaws interact with fewer neighbors than unpaired birds, flap their wings more slowly, which may save energy and flocks with more pairs exhibit shorter correlation length, which may lead to decreased group-level benefits (Ling et al., 2019a).

For the mammalian brain, already a cortical column with ∼10,000 neurons across its 6 layers provides a major modeling challenge with its diversity in cell types, cell connectivities, cellular and subcellular dynamics (Markram et al., 2015;Dura-Bernal et al., 2019). The type of dynamics that in principle can be generated in these high-dimensional models is not easily constrained and can range from large-scale synchronized oscillations to more local, sometimes sequential activity. With respect to the latter and in analogy to how the social relationships affect correlations in jackdaw flocks (Ling et al., 2019a), it has been shown that some neurons (leaders) consistently fire earlier than others in spontaneous bursts of activity *in vitro* (Eytan and Marom, 2006;Eckmann et al., 2008;Orlandi et al., 2013;Pasquale et al., 2017). Yet, it is currently not known how the heterogeneity of cell types, layers and areas contribute to scale-free correlation lengths measured in the awake brain at macro-, meso- and microscale. In a first attempt to address this issue, we studied functional subnetworks in cortical circuits, such as the one formed by orientation selective, i.e. “tuned” cells with similar tuning preference in V1 (Palagina et al., 2019). When separately analyzing tuned and non-tuned cells, despite significant changes in the absolute value of correlation changes (evidencing the different structure present in these subgroups), we were able to show that scale-free correlations are present along the tuning dimension (Ribeiro et al., 2020). We note that the subgroup not included in the analysis was still participating in the overall network response and that this finding does not exclude the possibility that both subgroups are essential to create the observed scale-free correlations for both subnetworks.

### Nature of the phase transition underlying the collective properties of animal groups and neuronal populations

A variety of collective states can be observed in animal groups. For instance, Tunstrøm and colleagues have shown that golden shiner schools can present three dynamically-stable collective states, namely swarm, polarized and milling, with frequent transitions between them (Tunstrøm et al., 2013). Naturally, different types of collective states and accompanied transitions between those states are necessary for different animal species. Here we discussed coordination or synchronization in animal groups in the context of emerging of directional order (or onset of collective motion) (Vicsek and Zafeiris, 2012) and condensation or clustering transitions (Chen et al., 2012;Calvao and Brigatti, 2019). Modeling work reflects this wide variety in collective behavior, which have been linked to different types of phase transitions, mainly of the discontinuous type (first order transitions, including hysteresis and metastability; e.g. (Couzin et al., 2002;Chate et al., 2008;Hein et al., 2015;Calvao and Brigatti, 2019)) or the continuous type (second order, in line with criticality; e.g. (Barberis and Albano, 2014;Calovi et al., 2015;Feinerman et al., 2018)), or both (Huepe et al., 2011). Even within one model, the type of phase transition encountered can be sensitive to the specific model parameters and simulations conducted. For example, the original introduction of the Vicsek model (Vicsek et al., 1995) suggested a second-order phase transition, yet, clear discontinuities where identified particularly when adding aggregational terms and/or allowing noise to be directly added to the neighborhood computation (Gregoire and Chate, 2004;Chate et al., 2008). Finite-size effects can smooth a discontinuous transition making it appear continuous (Gregoire and Chate, 2004;Solon et al., 2015;Brown et al., 2020), yet within certain parameter regimes, the Vicsek model exhibits robust continuous transitions (Barberis and Albano, 2014). These commonly encountered sensitivities of abstract models to parameter regime and seemingly innocent model variation, necessarily call for elaborate experimental designs to validate models. For example, cooperative transport in ants was found to be more in line with a continuous phase transition when quantifying transport velocity for food pellets of different sizes (Feinerman et al., 2018).

The plethora of models that can be construed for brain networks ranging from abstract, binary neurons with random connectivity to detailed compartmental neuronal networks requires a prudent and stepwise alignment of theory and models with continuously improving experimental evidence. Here, we would like to point out the experimental demonstration of scale-free neuronal avalanches in isolated brain preparations in line with predictions for a critical branching process (Beggs and Plenz, 2003). This experimental finding suggested that system wide correlations form spontaneously in a fluctuation dominated brain state, with low and sparse rate. The experimental demonstration of scale-free (most-often weak) correlations for spontaneous and evoked neuronal activity in the awake brain in the presence of scale-invariant neuronal avalanches has been reliably found at the macroscale (Expert et al., 2011;Fraiman and Chialvo, 2012;Tagliazucchi et al., 2012), meso and microscale (Ribeiro et al., 2020). Importantly, LFP avalanches that show scale-free correlations also exhibit a scaling collapse with an exponent of 2 for mean size vs. duration and an inverted parabolic profile in line with prediction for critical dynamics (Miller et al., 2019). It is this body of experimental results in the awake cortex (Scott et al., 2014;Bellay et al., 2015), which forms the seed for a more comprehensive understanding of the mechanisms ruling the scale-free dynamics in brain activity.

A variety of alternative models and simulations often exhibit significant shortcomings when accounting for the above-mentioned body of experimental findings. For example, the identification of universality classes that deviate from the directed percolation model have been found to be indecisive to explain neural data obtained from the anesthetized or sleep state under severe subsampling conditions (Fontenele et al., 2019;Carvalho et al., 2020). Similarly, neuronal models that feature a first order transition between a low and high activity mode switched randomly by external noise (Scarpetta and de Candia, 2013;Scarpetta et al., 2018) do not exhibit the experimentally reported scaling relation between size and duration of avalanches, but instead exhibit a scaling in line with a memory-less noise process (Villegas et al., 2019). Shortcomings can also be identified in the Landau-Ginzburg scenario introduced recently to simulate avalanches in neuronal networks (di Santo et al., 2016;Buendia et al., 2020) which, under certain parameter values, exhibits a first order transition, hysteresis and exponents similar to those of a critical branching process, yet the temporal avalanche profile differs from the inverted-parabola identified experimentally. In addition, the disorder-synchronization phase transition in that model gives rise to statistically distinct giant (“king”) avalanches found only in disinhibited brain activity similar to epileptic seizures.

As a final reflection on this aspect, it needs to be noted that in contrast with the empirical solitude of the finding of neuronal avalanches a decade and half ago, the field is currently populated by a large variety of not-always self-consistent models. It seems that a fruitful avenue now might be to balance the modeling efforts with a careful analysis of the continuously improving sophisticated experimental evidence at hand.

### Importance of scale-free correlations for brain function

A large body of modeling work and some experimental evidence have shown that scale-free correlations are beneficial, providing key advantages to animals living in groups. For example, Rauch et al. (1995) showed the emergence of self-organized trails near a critical density of foraging ants. The length of these trails exceeded several orders of magnitude the ants perceptual scale, being another example of long-range correlations. In the same line, it has been shown that evolutionary pressure could move fish schools toward an optimized state near a discontinuous phase transition in an evolutionary model, where local environmental perturbations can cause changes in the collective school state (Hein et al., 2015). Using the Vicsek model for flocks (Vicsek et al., 1995), it has been shown that information transmission is maximized near the phase transition (Fig. 2A; (Vanni et al., 2011;Lukovic et al., 2014)), which as discussed in the previous subsection could have an underlying first-order origin. The enlarged correlations arising as a result from this maximized information transmission, lead to optimized response to predators (Mateo et al., 2017), in line with what has been observed in data-driven models of fish schools (Calovi et al., 2015) or sheep herds (Ginelli et al., 2015) near criticality. It has also been shown that the efficiency of computations in the Grégoire and Chaté model (2004) is maximized at the phase transition (Crosato et al., 2018).

**Figure 2:**
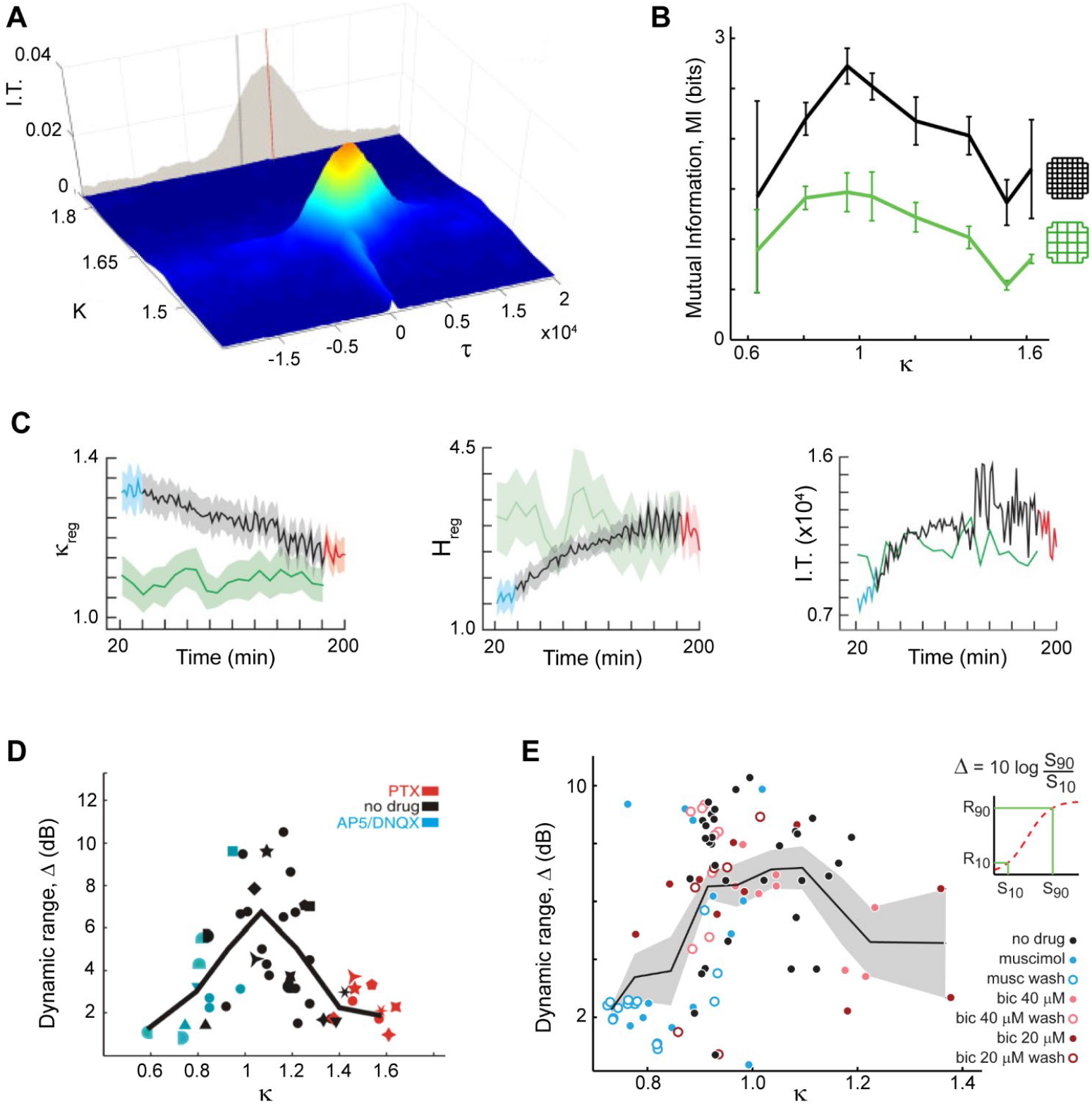
Criticality and scale-free organization provide key advantages for both flocks and brains such as maximal information transmission and dynamic range. **(A)**In a decision-making model of flock behavior, information transmission peaks at criticality, shown by a peak in mutual information for K ∼ 1.62 and τ ∼ 2500 (critical point). Adapted from (Lukovic et al., 2014). **(B)** In organotypic cultures grown from rodent brains, information transmission peaks when spontaneous neuronal activity displays scale-free neuronal avalanches, in line with expectation from criticality (κ ∼ 1). Two different coarse-graining levels are shown (*colors*). Proximity to criticality, i.e. scale-free avalanches, is controlled through pharmacological manipulation of the cultures. Adapted from (Shew et al., 2011). **(C)** Information transmission is maximized as mice recover from anesthesia, establishing neuronal avalanches. *Left*: Criticality distance measure approaches 1 (critical point) as time from anesthesia application (in min) passes. Anesthetized (*blue*), recently awake (*red*) and fully awake (*green*) states are highlighted. Entropy (*middle*) and information transmission (*right*) reaches a maximum as mice recover from anesthesia and reestablishing neuronal avalanches. Adapted from (Fagerholm et al., 2016). **(D)** In organotypic cortex cultures, the dynamic range peaks when neuronal avalanches emerge and can be reduced when pharmacologically changing the natural excitation/inhibition balance. Adapted from (Shew et al., 2009). **(E)** Using microelectrode array recordings in rats in vivo, the peak of dynamic range was demonstrated using natural stimuli and changes in excitation/inhibition balance through local pharmacological manipulation. Adapted from (Gautam et al., 2015).

On the brain side, theory and model simulations on critical dynamics in neuronal networks has proposed many advantages in information processing, some of which have been demonstrated experimentally, specifically when using pharmacological manipulations to move cortical networks away from neuronal avalanche dynamics (Fig. 2B – E; for reviews, see e.g. (Shew and Plenz, 2013;Cocchi et al., 2017)). For example, the dynamic range, which measures the range of stimulus intensity a network is able to differentiate, has been proposed to maximize at criticality by Kinouchi and Copelli (2006) and was demonstrated experimentally (Figs. 2D, E; (Shew et al., 2009;Gautam et al., 2015)).

Another parallel between scale-free flocks and brains is the presence of decentralized signal processing. This aspect has gained increased attention in the context of artificial intelligence, with many studies proposing the usage of artificial swarm systems (Hornischer et al., 2019;Sueoka et al., 2019). The brain also provides inspiration for these systems: Monaco and colleagues (Monaco et al., 2019) proposed an analogy between these multi-agent robotic platforms and place cells in the hippocampus, suggesting improvements to current models that follow solutions found by brain circuits. Startle responses in animal populations can trigger escape waves (Herbert-Read et al., 2015;Sosna et al., 2019), in the latter case yielding heavy-tail cascade size distributions and involve distributed repositioning of in the swarm beyond an individual’s sensitivity changes to perturbation. The initiation and spread of such local response bears similarities to branching process dynamics suggesting promising similarities with critical brain dynamics.

## Conclusions

The emergence of order in systems composed of a myriad of small entities exhibits many parallels between animal groups and neuronal populations in the brain. We summarized new experimental findings for the brain on the emergence of scale-invariant correlations and scale-invariant population sizes and discussed their similarities and differences compared to collective behavior in animals. We show that for both fields of research there are fascinating arguments for systems to be positioned near a phase transition to support propagation of local information throughout the entire system. Future experimental work on the role of cell types and microcircuit mechanisms in maintaining these scale- free dynamical features are crucial for understanding how the brain processes information.

## Author Contributions

All authors have made a substantial, direct and intellectual contribution to the work, and approved it for publication.

## Acknowledgments

We thank members of the Plenz lab for discussions.

## References

Aoki, I. (1982). A Simulation Study on the Schooling Mechanism in Fish. Bulletin of the Japanese Society of Scientific Fisheries 48, 1081–1088.

Arieli, A., Sterkin, A., Grinvald, A., and Aertsen, A. (1996). Dynamics of ongoing activity: Explanation of the large variability in evoked cortical responses. Science 273, 1868–1871.

Attanasi, A., Cavagna, A., Del Castello, L., Giardina, I., Melillo, S., Parisi, L., Pohl, O., Rossaro, B., Shen, E., Silvestri, E., and Viale, M. (2014a). Collective Behaviour without Collective Order in Wild Swarms of Midges. Plos Computational Biology 10.

Attanasi, A., Cavagna, A., Del Castello, L., Giardina, I., Melillo, S., Parisi, L., Pohl, O., Rossaro, B., Shen, E., Silvestri, E., and Viale, M. (2014b). Finite-Size Scaling as a Way to Probe Near-Criticality in Natural Swarms. Physical Review Letters 113.

Bak, P. (1996). How Nature Works. New York: Copernicus.

Ballerini, M., Calbibbo, N., Candeleir, R., Cavagna, A., Cisbani, E., Giardina, I., Lecomte, V., Orlandi, A., Parisi, G., Procaccini, A., Viale, M., and Zdravkovic, V. (2008). Interaction ruling animal collective behavior depends on topological rather than metric distance: Evidence from a field study. Proceedings of the National Academy of Sciences of the United States of America 105, 1232–1237.

Barberis, L., and Albano, E.V. (2014). Evidence of a robust universality class in the critical behavior of self-propelled agents: Metric versus topological interactions. Physical Review E 89.

Bednekoff, P.A., and Lima, S.L. (1998). Randomness, chaos and confusion in the study of antipredator vigilance. Trends in Ecology & Evolution 13, 284–287.

Beggs, J.M., and Plenz, D. (2003). Neuronal avalanches in neocortical circuits. Journal of Neuroscience 23, 11167–11177.

Belden, J., Mansoor, M.M., Hellum, A., Rahman, S.R., Meyer, A., Pease, C., Pacheco, J., Koziol, S., and Truscott, T.T. (2019). How vision governs the collective behaviour of dense cycling pelotons. Journal of the Royal Society Interface 16.

Bellay, T., Klaus, A., Seshadri, S., and Plenz, D. (2015). Irregular spiking of pyramidal neurons organizes as scale-invariant neuronal avalanches in the awake state. eLife 4, e07224.

Bialek, W., Cavagna, A., Giardina, I., Mora, T., Silvestri, E., Viale, M., and Walczak, A.M. (2012). Statistical mechanics for natural flocks of birds. Proceedings of the National Academy of Sciences of the United States of America 109, 4786–4791.

Boucsein, C., Nawrot, M.P., Schnepel, P., and Aertsen, A. (2011). Beyond the cortical column: abundance and physiology of horizontal connections imply a strong role for inputs from the surround. Front Neurosci 5, 32.

Brown, J., Bossomaier, T., and Barnett, L. (2020). Information flow in finite flocks. Scientific Reports 10.

Bruno, R.M., and Sakmann, B. (2006). Cortex Is Driven by Weak but Synchronously Active Thalamocortical Synapses. Science 312, 1622.

Buendia, V., Di Santo, S., Villegas, P., Burioni, R., and Munoz, M.A. (2020). Self-organized bistability and its possible relevance for brain dynamics. Physical Review Research 2, 013318.

Calovi, D.S., Lopez, U., Schuhmacher, P., Chate, H., Sire, C., and Theraulaz, G. (2015). Collective response to perturbations in a data-driven fish school model. Journal of the Royal Society Interface 12.

Calvao, A.M., and Brigatti, E. (2019). Collective movement in alarmed animals groups: A simple model with positional forces and a limited attention field. Physica a-Statistical Mechanics and Its Applications 520, 450–457.

Carvalho, T.T., Fontenele, A.J., Girardi-Schappo, M., Feliciano, T., Aguiar, L.A., Silva, T.P., De Vasconcelos, N.A., Carelli, P.V., and Copelli, M. (2020). Subsampled directed-percolation models explain scaling relations experimentally observed in the brain. arXiv preprint arXiv:2007.13813.

Cavagna, A., Cimarelli, A., Giardina, I., Parisi, G., Santagati, R., Stefanini, F., and Viale, M. (2010). Scale-free correlations in starling flocks. Proceedings of the National Academy of Sciences of the United States of America 107, 11865–11870.

Cavagna, A., Del Castello, L., Dey, S., Giardina, I., Melillo, S., Parisi, L., and Viale, M. (2015). Short-range interactions versus long-range correlations in bird flocks. Physical Review E 92.

Cavagna, A., Giardina, I., and Grigera, T.S. (2018). The physics of flocking: Correlation as a compass from experiments to theory. Physics Reports-Review Section of Physics Letters 728, 1–62.

Chate, H., Ginelli, F., Gregoire, G., and Raynaud, F. (2008). Collective motion of self-propelled particles interacting without cohesion. Phys Rev E Stat Nonlin Soft Matter Phys 77, 046113.

Chen, X., Dong, X., Be’er, A., Swinney, H.L., and Zhang, H.P. (2012). Scale-Invariant Correlations in Dynamic Bacterial Clusters. Physical Review Letters 108.

Chialvo, D.R. (2010). Emergent complex neural dynamics. Nature Physics 6, 744–750.

Cocchi, L., Gollo, L.L., Zalesky, A., and Breakspear, M. (2017). Criticality in the brain: A synthesis of neurobiology, models and cognition. Progress in Neurobiology 158, 132–152.

Couzin, I.D., Krause, J., James, R., Ruxton, G.D., and Franks, N.R. (2002). Collective memory and spatial sorting in animal groups. Journal of Theoretical Biology 218, 1–11.

Crosato, E., Spinney, R.E., Nigmatullin, R., Lizier, J.T., and Prokopenko, M. (2018). Thermodynamics and computation during collective motion near criticality. Physical Review E 97.

Cucker, F., and Smale, S. (2007). Emergent behavior in flocks. Ieee Transactions on Automatic Control 52, 852–862.

Di Santo, S., Burioni, R., Vezzani, A., and Munoz, M.A. (2016). Self-Organized Bistability Associated with First-Order Phase Transitions. Physical Review Letters 116.

Dura-Bernal, S., Suter, B.A., Gleeson, P., Cantarelli, M., Quintana, A., Rodriguez, F., Kedziora, D.J., Chadderdon, G.L., Kerr, C.C., Neymotin, S.A., Mcdougal, R.A., Hines, M., Shepherd, G.M.G., and Lytton, W.W. (2019). NetPyNE, a tool for data-driven multiscale modeling of brain circuits. eLife 8, e44494.

Eckmann, J.P., Jacobi, S., Marom, S., Moses, E., and Zbinden, C. (2008). Leader neurons in population bursts of 2D living neural networks. New Journal of Physics 10.

Ero, C., Gewaltig, M.O., Keller, D., and Markram, H. (2018). A Cell Atlas for the Mouse Brain. Frontiers in Neuroinformatics 12.

Evans, M.H.R., Lihou, K.L., and Rands, S.A. (2018). Black-headed gulls synchronise their activity with their nearest neighbours. Scientific Reports 8.

Expert, P., Lambiotte, R., Chialvo, D.R., Christensen, K., Jensen, H.J., Sharp, D.J., and Turkheimer, F. (2011). Self-similar correlation function in brain resting-state functional magnetic resonance imaging. J R Soc Interface 8, 472–479.

Eytan, D., and Marom, S. (2006). Dynamics and effective topology underlying synchronization in networks of cortical neurons. Journal of Neuroscience 26, 8465–8476.

Fagerholm, E.D., Scott, G., Shew, W.L., Song, C.C., Leech, R., Knopfel, T., and Sharp, D.J. (2016). Cortical Entropy, Mutual Information and Scale-Free Dynamics in Waking Mice. Cerebral Cortex 26, 3945–3952.

Feinerman, O., Pinkoviezky, I., Gelblum, A., Fonio, E., and Gov, N.S. (2018). The physics of cooperative transport in groups of ants. Nature Physics 14, 683–693.

Fontenele, A.J., De Vasconcelos, N.a.P., Feliciano, T., Aguiar, L.a.A., Soares-Cunha, C., Coimbra, B., Dalla Porta, L., Ribeiro, S., Rodrigues, A.J., Sousa, N., Carelli, P.V., and Copelli, M. (2019). Criticality between cortical states. Physical Review Letters 122, 208101.

Fraiman, D., and Chialvo, D.R. (2012). What kind of noise is brain noise: anomalous scaling behavior of the resting brain activity fluctuations. Frontiers in Physiology 3.

Gautam, H., Hoang, T.T., Mcclanahan, K., Grady, S.K., and Shew, W.L. (2015). Maximizing Sensory Dynamic Range by Tuning the Cortical State to Criticality. Plos Computational Biology 11.

Gelblum, A., Pinkoviezky, I., Fonio, E., Gov, N.S., and Feinerman, O. (2016). Emergent oscillations assist obstacle negotiation during ant cooperative transport. Proceedings of the National Academy of Sciences of the United States of America 113, 14615–14620.

Ginelli, F., and Chate, H. (2010). Relevance of Metric-Free Interactions in Flocking Phenomena. Physical Review Letters 105.

Ginelli, F., Peruani, F., Pillot, M.H., Chate, H., Theraulaz, G., and Bon, R. (2015). Intermittent collective dynamics emerge from conflicting imperatives in sheep herds. Proceedings of the National Academy of Sciences of the United States of America 112, 12729–12734.

Goldstone, J. (1961). Field Theories with Superconductor Solutions. Nuovo Cimento 19, 154–164.

Gouwens, N.W., Sorensen, S.A., Berg, J., Lee, C., Jarsky, T., Ting, J., Sunkin, S.M., Feng, D., Anastassiou, C.A., Barkan, E., Bickley, K., Blesie, N., Braun, T., Brouner, K., Budzillo, A., Caldejon, S., Casper, T., Castelli, D., Chong, P., Crichton, K., Cuhaciyan, C., Daigle, T.L., Dailey, R., Dee, N., Desta, T., Ding, S.L., Dingman, S., Doperalski, A., Dotson, N., Egdorf, T., Fisher, M., De Frates, R.A., Garren, E., Garwood, M., Gary, A., Gaudreault, N., Godfrey, K., Gorham, M., Gu, H., Habel, C., Hadley, K., Harrington, J., Harris, J.A., Henry, A., Hill, D., Josephsen, S., Kebede, S., Kim, L., Kroll, M., Lee, B., Lemon, T., Link, K.E., Liu, X.X., Long, B., Mann, R., Mcgraw, M., Mihalas, S., Mukora, A., Murphy, G.J., Ng, L., Ngo, K., Nguyen, T.N., Nicovich, P.R., Oldre, A., Park, D., Parry, S., Perkins, J., Potekhina, L., Reid, D., Robertson, M., Sandman, D., Schroedter, M., Slaughterbeck, C., Soler-Llavina, G., Sulc, J., Szafer, A., Tasic, B., Taskin, N., Teeter, C., Thatra, N., Tung, H., Wakeman, W., Williams, G., Young, R., Zhou, Z., Farrell, C., Peng, H.C., Hawrylycz, M.J., Lein, E., Ng, L., Arkhipov, A., Bernard, A., Phillips, J.W., Zeng, H.K., and Koch, C. (2019). Classification of electrophysiological and morphological neuron types in the mouse visual cortex. Nature Neuroscience 22, 1182-+.

Greenberg, R. (2000). “Birds of many feathers: the formation and structure of mixed species flocks of forest birds,” in On the move: how and why animals travel in groups, eds. S. Boinski & P. Garber. (Chicago: Chicago University Press).

Gregoire, G., and Chate, H. (2004). Onset of collective and cohesive motion. Phys Rev Lett 92, 025702.

Gregoire, G., Chate, H., and Tu, Y.H. (2003). Moving and staying together without a leader. Physica D-Nonlinear Phenomena 181, 157–170.

Haimovici, A., Tagliazucchi, E., Balenzuela, P., and Chialvo, D.R. (2013). Brain Organization into Resting State Networks Emerges at Criticality on a Model of the Human Connectome. Physical Review Letters 110.

Hamilton, W.D. (1971). Geometry for Selfish Herd. Journal of Theoretical Biology 31, 295-&.

Hein, A.M., Rosenthal, S.B., Hagstrom, G.I., Berdahl, A., Torney, C.J., and Couzin, I.D. (2015). The evolution of distributed sensing and collective computation in animal populations. Elife 4.

Herbert-Read, J.E., Buhl, J., Hu, F., Ward, A.J.W., and Sumpter, D.J.T. (2015). Initiation and spread of escape waves within animal groups. Royal Society Open Science 2.

Herbert-Read, J.E., Perna, A., Mann, R.P., Schaerf, T.M., Sumpter, D.J.T., and Ward, A.J.W. (2011). Inferring the rules of interaction of shoaling fish. Proceedings of the National Academy of Sciences of the United States of America 108, 18726–18731.

Hesse, J., and Gross, T. (2014). Self-organized criticality as a fundamental property of neural systems. Frontiers in Systems Neuroscience 8.

Hornischer, H., Herminghaus, S., and Mazza, M.G. (2019). Structural transition in the collective behavior of cognitive agents. Scientific Reports 9.

Huepe, C., Zschaler, G., Do, A.L., and Gross, T. (2011). Adaptive-network models of swarm dynamics. New Journal of Physics 13.

Huth, A., and Wissel, C. (1992). The Simulation of the Movement of Fish Schools. Journal of Theoretical Biology 156, 365–385.

Kertesz, J., and Wolf, D.E. (1989). Anomalous roughening in growth processes. Phys Rev Lett 62, 2571–2574.

King, A.J., Fehlmann, G., Biro, D., Ward, A.J., and Furtbauer, I. (2018). Re-wilding Collective Behaviour: An Ecolocical Perspective. Trends in Ecology & Evolution 33, 347–357.

Kinouchi, O., and Copelli, M. (2006). Optimal dynamical range of excitable networks at criticality. Nature Physics 2, 348–352.

Krause, J., and Ruxton, G.D. (2002). Living in Groups. Oxford: Oxford University Press.

Krebs, J.R. (1973). Social-Learning and Significance of Mixed-Species Flocks of Chickadees (Parus Spp). Canadian Journal of Zoology-Revue Canadienne De Zoologie 51, 1275–1288.

Levy, R.B., and Reyes, A.D. (2012). Spatial Profile of Excitatory and Inhibitory Synaptic Connectivity in Mouse Primary Auditory Cortex. Journal of Neuroscience 32, 5609–5619.

Ling, H.J., Mcivor, G.E., Van Der Vaart, K., Vaughan, R.T., Thornton, A., and Ouellette, N.T. (2019a). Costs and benefits of social relationships in the collective motion of bird flocks. Nature Ecology & Evolution 3, 943–948.

Ling, H.J., Mcivor, G.E., Van Der Vaart, K., Vaughan, R.T., Thornton, A., and Ouellette, N.T. (2019b). Local interactions and their group-level consequences in flocking jackdaws. Proceedings of the Royal Society B-Biological Sciences 286.

Lukovic, M., Vanni, F., Svenkeson, A., and Grigolini, P. (2014). Transmission of information at criticality. Physica a-Statistical Mechanics and Its Applications 416, 430–438.

Ma, S.-K. (1976). Modern Theory of Critical Phenomena. Reading, MA: Benjamin. Ma, S.-K. (1985). Statistical Mechanics. Philadelphia: World Scientific.

Markram, H., Muller, E., Ramaswamy, S., Reimann Michael w., Abdellah, M., Sanchez Carlos a., Ailamaki, A., Alonso-Nanclares, L., Antille, N., Arsever, S., Kahou Guy antoine a., BergerThomas k., Bilgili, A., Buncic, N., Chalimourda, A., Chindemi, G., Courcol, J.-D., Delalondre, F., Delattre, V., Druckmann, S., Dumusc, R., Dynes, J., Eilemann, S., Gal, E., Gevaert Michael e., Ghobril, J.-P., Gidon, A., Graham, Joe w., Gupta, A., Haenel, V., Hay, E., Heinis, T., Hernando, Juan b., Hines, M., Kanari, L., Keller, D., Kenyon, J., Khazen, G., Kim, Y., King James g., Kisvarday, Z., Kumbhar, P., Lasserre, S., Le bé, J.-V., Magalhães Brunor. C., Merchán-Pérez, A., Meystre, J., Morrice Benjamin r., Muller, J., Muñoz-Céspedes, A., Muralidhar, S., Muthurasa, K., Nachbaur, D.,, Newton Taylor h., Nolte, M., Ovcharenko, A., Palacios, J., Pastor, L., Perin, R., Ranjan, R., Riachi, I., Rodríguez, J.-R., Riquelme Juan l., Rössert, C., Sfyrakis, K., Shi, Y., Shillcock Julian c., Silberberg, G., Silva, R., Tauheed, F., Telefont, M., Toledo-Rodriguez, M., Tränkler, T., Van geit, W., Díaz Jafet v., Walker, R., Wang, Y., Zaninetta Stefano m., Defelipe, J., Hill Sean l., Segev, I., and Schürmann, F. (2015). Reconstruction and simulation of neocortical microcircuitry. Cell 163, 456–492.

Martys, N., Cieplak, M., and Robbins, M.O. (1991). Critical phenomena in fluid invasion of porous media. Phys Rev Lett 66, 1058–1061.

Mateo, D., Kuan, Y.K., and Bouffanais, R. (2017). Effect of Correlations in Swarms on Collective Response. Scientific Reports 7.

Meakin, P. (1987). “The Growth of Fractal Aggregates and their Fractal Measures,” in Phase Transitions and Critical Phenomena, eds. C. Domb & J.L. Lebowitz. (New York: Academic Press).

Miller, S.R., Yu, S., and Plenz, D. (2019). The scale-invariant, temporal profile of neuronal avalanches in relation to cortical γ–oscillations. Scientific Reports 9, 16403.

Millonas, M.M. (1992). A connectionist type model of self-organized foraging and emergent behavior in ant swarms. Journal of Theoretical Biology 159, 529–552.

Monaco, J.D., Hwang, G.M., Schultz, K.M., and Zhang, K.C. (2019). Cognitive swarming: An approach from the theoretical neuroscience of hippocampal function. Micro- and Nanotechnology Sensors, Systems, and Applications Xi 10982.

Mora, T., and Bialek, W. (2011). Are Biological Systems Poised at Criticality? Journal of Statistical Physics 144, 268–302.

Munn, C.A., and Terborgh, J.W. (1979). Multi-Species Territoriality in Neotropical Foraging Flocks. Condor 81, 338–347.

Nagy, M., Couzin, I.D., Fiedler, W., Wikelski, M., and Flack, A. (2018). Synchronization, coordination and collective sensing during thermalling flight of freely migrating white storks. Philosophical Transactions of the Royal Society B-Biological Sciences 373.

Orlandi, J.G., Soriano, J., Alvarez-Lacalle, E., Teller, S., and Casademunt, J. (2013). Noise focusing and the emergence of coherent activity in neuronal cultures. Nature Physics 9, 582–590.

Palagina, G., Meyer, J.F., and Smirnakis, S.M. (2019). Inhibitory Units: An Organizing Nidus for Feature-Selective SubNetworks in Area V1. Journal of Neuroscience 39, 4931–4944.

Pasquale, V., Martinoia, S., and Chiappalone, M. (2017). Stimulation triggers endogenous activity patterns in cultured cortical networks. Scientific Reports 7.

Plenz, D. (2012). Neuronal avalanches and coherence potentials. European Physical Journal-Special Topics 205, 259–301.

Plenz, D., and Niebur, E. (2014). Criticality in Neural Systems. Berlin: Wiley-VCH.

Powell, G.V.N. (1985). Sociobiology and adaptive significance of interspecific foraging flocks in the Neotropics. Ornithological Monographs 36, 713–732.

Rands, S.A., Muir, H., and Terry, N.L. (2014). Red deer synchronise their activity with close neighbours. Peerj 2.

Rauch, E.M., Millonas, M.M., and Chialvo, D.R. (1995). Pattern-formation and functionality in swarm models. Physics Letters A 207, 185–193.

Reynolds, C.W. (1987). Flocks, herds and schools: a distributed behavioral model. Computer Graphics 21, 25–34.

Ribeiro, T.L., Yu, S., Martin, D.A., Winkowski, D., Kanold, P., Chialvo, D.R., and Plenz, D. (2020). Trial-by-trial variability in cortical responses exhibits scaling in spatial correlations predicted from critical dynamics. submitted to bioRxiv.

Romanczuk, P., Couzin, I.D., and Schimansky-Geier, L. (2009). Collective Motion due to Individual Escape and Pursuit Response. Physical Review Letters 102.

Romenskyy, M., Herbert-Read, J.E., Ward, A.J.W., and Sumpter, D.J.T. (2017). Body size affects the strength of social interactions and spatial organization of a schooling fish (Pseudomugil signifer). Royal Society Open Science 4.

Scarpetta, S., Apicella, I., Minati, L., and De Candia, A. (2018). Hysteresis, neural avalanches, and critical behavior near a first-order transition of a spiking neural network. Physical Review E 97.

Scarpetta, S., and De Candia, A. (2013). Neural Avalanches at the Critical Point between Replay and Non-Replay of Spatiotemporal Patterns. Plos One 8.

Scott, G., Fagerholm, E.D., Mutoh, H., Leech, R., Sharp, D.J., Shew, W.L., and Knopfel, T. (2014). Voltage imaging of waking mouse cortex reveals emergence of critical neuronal dynamics. J. Neurosci. 34, 16611–16620.

Seeman, S.C., Campagnola, L., Davoudian, P.A., Hoggarth, A., Hage, T.A., Bosma-Moody, A., Baker, C.A., Lee, J.H., Mihalas, S., Teeter, C., Ko, A.L., Ojemann, J.G., Gwinn, R.P., Silbergeld, D.L., Cobbs, C., Phillips, J., Lein, E., Murphy, G., Koch, C., Zeng, H.K., and Jarsky, T. (2018). Sparse recurrent excitatory connectivity in the microcircuit of the adult mouse and human cortex. Elife 7.

Shew, W.L., and Plenz, D. (2013). The Functional Benefits of Criticality in the Cortex. Neuroscientist 19, 88–100.

Shew, W.L., Yang, H., Yu, S., Roy, R., and Plenz, D. (2011). Information capacity and transmission are maximized in balanced cortical networks with neuronal avalanches. J Neurosci 31, 55–63.

Shew, W.L., Yang, H.D., Petermann, T., Roy, R., and Plenz, D. (2009). Neuronal Avalanches Imply Maximum Dynamic Range in Cortical Networks at Criticality. Journal of Neuroscience 29, 15595–15600.

Solon, A.P., Chate, H., and Tailleur, J. (2015). From Phase to Microphase Separation in Flocking Models: The Essential Role of Nonequilibrium Fluctuations. Physical Review Letters 114.

Sosna, M.M.G., Twomey, C.R., Bak-Coleman, J., Poel, W., Daniels, B.C., Romanczuk, P., and Couzin, I.D. (2019). Individual and collective encoding of risk in animal groups. Proceedings of the National Academy of Sciences of the United States of America 116, 20556–20561.

Stanley, H.E. (1971). Introduction to phase transitions and critical phenomena. New York: Oxford University Press.

Strandburg-Peshkin, A., Twomey, C.R., Bode, N.W.F., Kao, A.B., Katz, Y., Ioannou, C.C., Rosenthal, S.B., Torney, C.J., Wu, H.S., Levin, S.A., and Couzin, I.D. (2013). Visual sensory networks and effective information transfer in animal groups. Current Biology 23, R709–R711.

Strombom, D. (2011). Collective motion from local attraction. Journal of Theoretical Biology 283, 145–151.

Sueoka, Y., Sato, Y., Ishitani, M., and Osuka, K. (2019). Analysis of push-forward model for swarm-like collective motions. Artificial Life and Robotics 24, 460–470.

Sumpter, D.J.T. (2006). The principles of collective animal behaviour. Philosophical Transactions of the Royal Society B-Biological Sciences 361, 5–22.

Tagliazucchi, E., Balenzuela, P., Fraiman, D., and Chialvo, D.R. (2012). Criticality in large-scale brain fMRI dynamics unveiled by a novel point process analysis. Front. Physiol. 3.

Tang, Q.Y., Hatakeyama, T.S., and Kaneko, K. (2020). Functional sensitivity and mutational robustness of proteins. Physical Review Research 2, 033452.

Tang, Q.Y., Zhang, Y.Y., Wang, J., Wang, W., and Chialvo, D.R. (2017). Critical Fluctuations in the Native State of Proteins. Physical Review Letters 118.

Terborgh, J. (1990). Mixed Flocks and Polyspecific Associations - Costs and Benefits of Mixed Groups to Birds and Monkeys. American Journal of Primatology 21, 87–100.

Tunstrøm, K., Katz, Y., Ioannou, C.C., Huepe, C., Lutz, M.J., and Couzin, I.D. (2013). Collective States, Multistability and Transitional Behavior in Schooling Fish. Plos Computational Biology 9.

Vanni, F., Lukovic, M., and Grigolini, P. (2011). Criticality and Transmission of Information in a Swarm of Cooperative Units. Physical Review Letters 107.

Vicsek, T., Czirok, A., Ben-Jacob, E., Cohen, I.I., and Shochet, O. (1995). Novel type of phase transition in a system of self-driven particles. Phys Rev Lett 75, 1226–1229.

Vicsek, T., and Zafeiris, A. (2012). Collective motion. Physics Reports-Review Section of Physics Letters 517, 71–140.

Villegas, P., Di Santo, S., Burioni, R., and Muñoz, M.A. (2019). Time-series thresholding and the definition of avalanche size. Physical Review E 100.

Wang, X., and Lu, J.H. (2019). Collective Behaviors Through Social Interactions in Bird Flocks. Ieee Circuits and Systems Magazine 19, 6–22.

Wilson, K.G. (1979). Problems in Physics with Many Scales of Length. Scientific American 241, 158-&.

Yu, S., Ribeiro, T.L., Meisel, C., Chou, S., Mitz, A., Saunders, R., and Plenz, D. (2017). Maintained avalanche dynamics during task-induced changes of neuronal activity in nonhuman primates. Elife 6.

